# Tradeoffs and cultural diversity

**DOI:** 10.1101/263905

**Authors:** Omer Karin, Uri Alon

## Abstract

Culture is humanity’s prime adaptation. Which cultural traits contribute to adaptive value, and how they do so, is, however, unclear and debated. Here, we apply an approach from systems biology, known as Pareto task inference (ParTI), to bring a fresh perspective to these questions. ParTI considers systems that need to perform multiple tasks. No system can be optimal at all tasks at once, leading to a fundamental tradeoff. Such tradeoffs constrain evolution, because adaptive change for one task may be prevented because it compromises other tasks. These constraints result in specific polygon patterns in multivariate trait data. ParTI detects these polygons and their vertices, in order to infer the number of adaptive tasks and their nature. Here, we applied ParTI to two datasets of human cultural traits, on Austronesian cultures and modern hunter-gatherers, adjusting for phylogeny and spatial diffusion effects. We find that these independent datasets show the hallmarks of a tradeoff between the same three tasks: resource defense, resource competition, and mobility/exchange. Specific combinations of cultural traits are adaptive for each of these tasks. We thus suggest that part of the diversity of human cultural traits is constrained by tradeoffs between key tasks.

## Introduction

Culture has helped humans to adapt to diverse environments. Cultural traits vary widely, and the goal of the field of cultural evolution is to understand the processes that lead to this diversity (1). A central question is whether cultural traits are adaptive at the level of societies, and if so, how. A recent theory, by Boyd and Richerson (2,3), proposes that processes of cultural evolution lead to adaptedness of cultural traits. The theory posits that between-society variation in cultural traits is transmitted, inherited, and maintained by various mechanisms, which include biased social learning, coordination payoffs, punishment of deviant behaviors and society-level symbolic boundaries (4). Intergroup competition, in the form of natural selection, selective imitation, and selective migration, acts on this variation (2,5–9). Cultural traits that contribute to fitness in this sense tend to be selected (copied, maintained).

Testing such theories requires identifying which cultural traits are adaptive, and what is their function. This is difficult, since unlike biological traits such as physiological features (10) or gene expression (11), the societal-level function of cultural traits cannot be readily measured or manipulated in a controlled setting. Current methods mostly rely on correlations between traits and environmental features (5,12). It is therefore of interest to develop new methods to rigorously test whether cultural traits are adaptive, and, if so, what is their function.

We address this by applying an approach from systems biology to understand multi-task evolution in organisms, and adapt it to infer the society-level function of cultural traits directly from cross-cultural datasets. This method is applicable to the case in which performance at several tasks contributes to fitness. Thus, following the notion of performance introduced by Arnold for organic evolution (13), fitness as a function of traits 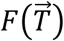 can be written in terms of performance functions *P*_1_, *P*_2_, … *P*_*k*_, one for each task, such that 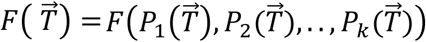, where *F* is an increasing function of the performances The idea is that each cultural trait may contribute to (or reduce) performance in each task. Because no single combination of traits can be optimal at all tasks at once, there is a fundamental trade-off.

A recent theory, called Pareto task inference (ParTI) (14,15), addresses this situation and provides a way to infer the number and identity of the tasks (Figure 1). ParTI theory shows that, under mild assumptions, the trait combinations that maximize *F* must fall inside a low-dimensional polygonal shape in trait space (15). Each vertex of the polygon is a set of traits 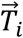 that is optimal for one of the tasks *i* (that is, the trait vector 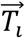 that maximizes one of the performance functions *P*_*i*_). Thus, the number of vertices is equal to the number of tasks: For two tasks, the data should fall on a line segment, whose ends are the two vertices; for three tasks, it should fall in a triangle with three vertices and so on. Specialists at each task lie near a vertex, and generalists lie in the center of the polygon. The intuitive reason for the polygon is that any point outside of the polygon has a point inside the polygon that is closer to all vertices, and hence has better performance at all tasks and hence higher fitness.

**Figure 1.**
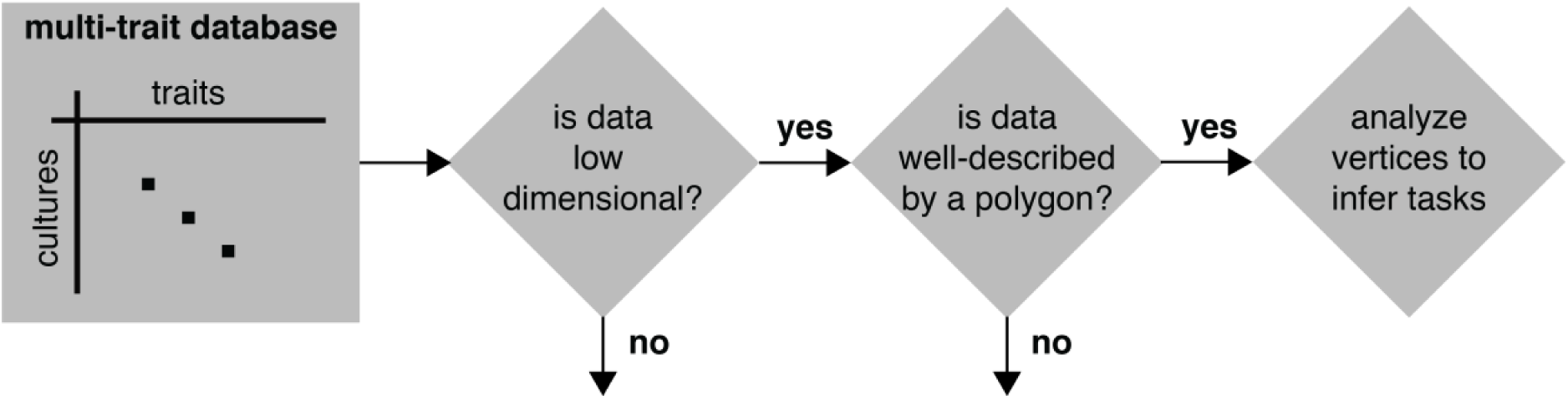
Schema of ParTI algorithm to infer tasks and trade-offs from multivariate trait data. ParTI receives as input a dataset where rows are cultures and columns are traits. ParTI tests the dimensionality of the dataset, and asks whether it is well-described by a polygon according to statistical tests. If the data indeed falls on a low-dimensional polygon, it infers its vertices. The number of vertices is equal to the number of tasks to be inferred. The cultures closest to each vertex are assumed to specialize in a certain task, and information about these cultures and the traits that make up the vertex is used to infer the task for each vertex.

ParTI algorithms determine the best-fit polygon (line, triangle, tetrahedron, etc.) that describes the data and its vertices. The algorithm uses statistical tests for the quality of the fit to the polygon, by comparing the data to shuffled data (Methods). It thus discovers the number of tasks, and, based on the cultures closest to each vertex, provides information that can be used to infer the tasks.

ParTI was used to define evolutionary tasks and tradeoffs in several biological domains, including gene expression in cells (16,17), development (18), animal morphology (19), and life-history strategies (20) (Supplementary Information 1). Due to the generality of the assumptions underlying ParTI, we reasoned that it might also apply to cultural evolution. In this case, the input data is multivariate cross-cultural trait datasets. Here we use ParTI to test hypothesis regarding adaptedness of cultural traits, and to infer the number and nature of the tasks that societies face, and which cultural traits are associated with these tasks.

## Methods

### Cross-cultural datasets

The *Pulotu* (21) dataset on traits of 113 Austronesian cultures, linked to a language-based family tree, was downloaded from https://pulotu.econ.mpg.de/. We used 46 traits that are cultural (i.e. not physical) and belong to the traditional time focus as defined by Watts et al. (21) (see Supplementary Information 2 for data preparation).

*Binford’s Hunter-Gatherer (BHG)* (22) dataset, that includes cultural traits for 339 modern hunter-gatherer cultures, was downloaded from http://ajohnson.sites.truman.edu/data-and-program/ (see Supplementary Information 3 for data preparation). The two datasets differ both in the mode of subsistence of societies that are included (*Pulotu* includes mostly agricultural societies) and in the cultural traits measured, which are largely non-overlapping between the datasets.

In Supplementary Tables 1, 2 we list the traits used for our analysis (codebooks for traits are available in the links above). Importantly, we verified that the results are not confounded by the structure of the dataset. That is, the empty regions in the trait-space of both datasets consist of (*a-priori*) feasible trait combinations (see Supplementary Information 5 for discussion).

### Language tree phylogenies

To generate a phylogenetic null model for principal component analysis and for polygon statistical significance, we used a tree with informative branch lengths for 93 Austronesian cultures from the *Pulotu* dataset (see Watts et al. (23)), `tree1`. For phylogenetic adjustment of enrichment calculations, which does not require informative branch lengths, we constructed a larger tree, `tree2`, by adding the other Austronesian cultures according to their language families (`tree2`, provided in Supplementary Information 8). Similarly, we constructed a phylogenetic tree for the cultures in *Binford’s Hunter-Gatherer* according to language families (`tree3`, provided in Supplementary Information 9).

### Number of significant principal components

To test the effective dimensionality of *Pulotu*, we determined the number of principal components that cannot be explained by a phylogenetic null model in which cultural traits are acquired and lost at random based on `tree1` (see Supplementary Information 4 for details).

### Polytope inference and statistical significance

To infer the vertices of the polytopes (polytopes are the generalization of polygons and polyhedra to any dimension) we applied Principal Convex Hull Analysis (PCHA) (14,24) as implemented in the python package *py_pcha*. Given an *n*-dimensional dataset and a number of vertices *k*, PCHA computes a convex hull with *k* (*n*-dimensional) vertices that best accounts for the variance in the data (14). This differs from PCA, which attempts to recover the orthogonal axes that explain the most variance, which need not correspond to the relevant trade-off space. We determined the number of vertices to test, k, by using the estimated dimension D of the data estimated by PCA, by *k*=*D*+*1*. Thus, two-dimensional data is fit to a triangle with three vertices, one-dimensional data to a line segment with two vertices, etc. We tested statistical significance of polygons by using the t-ratio test as specified by Shoval et al. (15). The t-ratio is the ratio between the areas (or volume) of the convex hull of the data and the area (or volume) of the minimal enclosing polygon. The tighter the polygon fits the data, the closer the t-ratio is to one. The idea is that a good fit of a polygon means that the external outline of the data cloud (its convex hull) has almost the same volume as the polygon (there is little empty space between the polygon and the data). A p-value was computed by the probability that the t-ratio of data shuffles is smaller or equal to the t-ratio of the real data based on 10,000 data shuffles (see Supplementary Information 5). In cases where we adjusted for phylogeny and spatial effects, the data was shuffled using null models hat preserve the phylogenetic and geographic structure (Supplementary Information 5).

### Trait regression according to distance from vertices

To infer tasks, we reasoned that traits relevant to a task should appear most frequently in the cultures closest to the vertex associated with that task. We tested this by performing regression, where the response variable is the trait and the predictor variable is the Euclidean distance from vertex in the space of the first two PCs. To adjust for spatial and phylogenetic effects, we added to each regression, and for each trait, covariates for the average trait value among the 10 nearest phylogenetic neighbors (`tree2` and `tree3`) and 10 nearest spatial neighbors (see Supplementary Information 6 for details), similar to the adjustment performed by Botero et al. (25). For binary variables, we performed logistic regression, and for other variables, we performed linear regression. We controlled for multiple hypotheses testing errors using FDR (Supplementary Information 6).

## Results

### ParTI analysis of the Pulotu dataset of Austronesian cultures

Using ParTI analysis, we find that the distribution of traits for the *Pulotu* cultures is well described by a triangle (p<10^-3^) (Figure 2A, Supplementary Information 5). We plot this triangle in the plane of the first two principal components, which explain 20% of the variance and are significant compared to a phylogenetic null model (Methods). The triangle is populated almost uniformly by cultures. Many a-priori possible combinations of traits do not seem to occur, resulting in the empty trait-space around the triangle. Shuffled data does not fill a triangle (Figure 2B), also when taking into account phylogeny (Supplementary Information 5).

**Figure 2.**
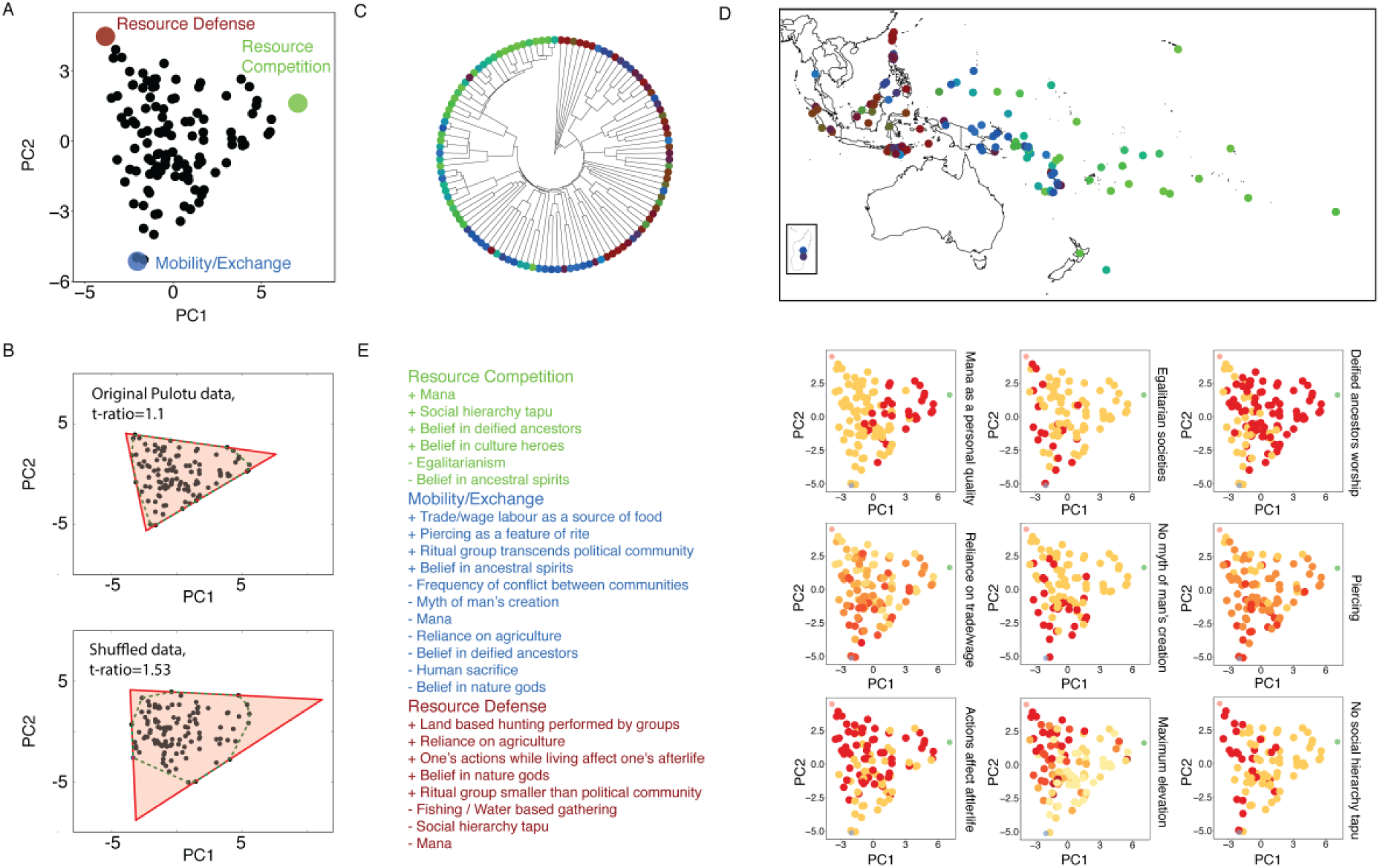
ParTI analysis of *Pulotu* dataset suggests three tasks. (A) *Pulotu* cultural traits are well described by a triangle (p<10^-3^) in the plain of the first two principal components. The three vertices of the triangle correspond to adaptive trait complexes for the putative tasks of mobility/exchange, resource defense and resource competition. (B) To test for statistical significance, we used the t-ratio test (15). The t-ratio is the ratio between the areas of the convex hull (pink) and the area of the minimal enclosing triangle. A t-ratio close to 1 means the data fills the triangle. Shuffled datasets, on the other hand, do not fill their enclosing triangles as tightly. (C) A consensus language tree for Austronesian languages. Cultures are color coded according to their relative distance from each vertex of the triangle. (D) A geographical map, with color coding as in C. (E) To infer tasks for each vertex, we seek traits that appear at higher frequency in the data points closest to the vertex. Such strongly enriched traits are listed. Sign (+/-) indicates positive or negative enrichment. Nine of the enriched traits are visualized on the data points, with red (yellow) corresponding to a high (low) level of the trait. For example, *no myth of mans’ creation* is found predominantly in cultures near the mobility/exchange vertex, whereas *deified ancestor worship* is frequent in cultures closest to the resource-competition vertex.

According to Pareto theory, this triangle indicates three tasks that have tradeoffs between them. The three vertices of the triangle are combinations of cultural traits that are potentially adaptive for these three tasks (see Table S1). Below, we infer the three putative tasks. We label the three vertices by these tasks – Resource Competition, Mobility/Exchange and Resource Defense - and describe how the tasks were inferred below.

In order to study the relation between the traits and language phylogeny, we displayed the cultures on a language tree. Cultures are color-coded according to their distances from the vertices of the triangle (Figure 2C). For example, cultures colored red are closest to the Resource Defense vertex on the triangle. We note an early-late phylogenetic gradient, from Resource Defense to Mobility/Exchange to Resource Competition. Similarly, we displayed the cultures on a geographical map (Figure 2D), showing an east-west geographical gradient from Resource Defense to Mobility/Exchange to Resource Competition. There are notable exceptions in which cultures that are close geographically and phylogenetically are not close on the triangle (have different color codes in Figure 2CD), suggesting that geographical proximity and phylogenetic inertia are not the only forces at play. Below, we adjusted for phylogenetic and geographic gradients in our analysis.

### Characterization of tasks for *Pulotu*

To characterize the underlying tasks, we asked which cultural traits characterize the cultures closest to each vertex. For this purpose, we used regression for each trait on its distance on the triangle from each vertex, adjusted for spatial and phylogenetic effects (Methods), correcting for multiple hypotheses testing errors. Each of the three vertices shows distinct cultural traits (Figure 2E).

The cultures closest to the Resource Competition vertex are the stratified Polynesian cultures (26,27). Closeness to this vertex is strongly associated with *mana* (supernatural power), especially when linked with descent and social status, and with the *worship of deified ancestors*. These traits are related to colonization and conquest - founders of colonies become revered ancestors and their descendants enjoy high status (“founder-focused ideology”) (26,28). Other traits found near this vertex include *low egalitarianism* and the *existence of a social hierarchy tapu*.

The Mobility/Exchange vertex is associated with cultures that live in areas of low resource density. The cultures closest to this vertex include the sea-nomad cultures of *Sama Dilaut* and *Moken* and the sea dwelling people of *Manus*. The Polynesian and Micronesian cultures that lie on low atolls are also close to this vertex (Supplementary Information 7). The Mobility/Exchange vertex is enriched with *infrequent conflict between communities, high reliance on trade and wage labor* rather than on agriculture, and *large ritual social groups* that transcend political communities. It is enriched with *piercing as a feature of rite*, as well as with *belief in ancestral spirits* (which have a smaller sphere of influence than do *deified ancestors*). Otherwise, this vertex is characterized by its lacks in religious complexity – *lack of mana, lack of human sacrifice, lack of a myth of man’s creation*, and *lack of belief in deified ancestors* and in *nature gods*.

The third vertex, Resource Defense, is enriched with highlander cultures that rely on *agriculture* and *land-based hunting* as opposed to fishing and water-based gathering, as well as with *belief in nature gods*, and with the *belief that the actions of oneself affect the nature of one’s afterlife*.

### ParTI analysis of the *BHG* dataset of modern hunter-gatherer cultures

To test the generality of these findings, we analyzed an independent dataset of modern hunter-gatherer cultures (22) (Figure 3). Considering the first two principal components, which explain 33% of the variance of this dataset, we again find a distribution inside a triangle (p<10^-3^) with three well-defined vertices (Figure 3A, Supplementary Information 5 and Table S2). For reasons discussed below, we assign the same three labels to these vertices. As with the *Pulotu* dataset, the traits show geographical trends (Figure 3B), and hence we performed regression adjusted for phylogenetic and spatial effects (Supplementary Information 6).

**Figure 3.**
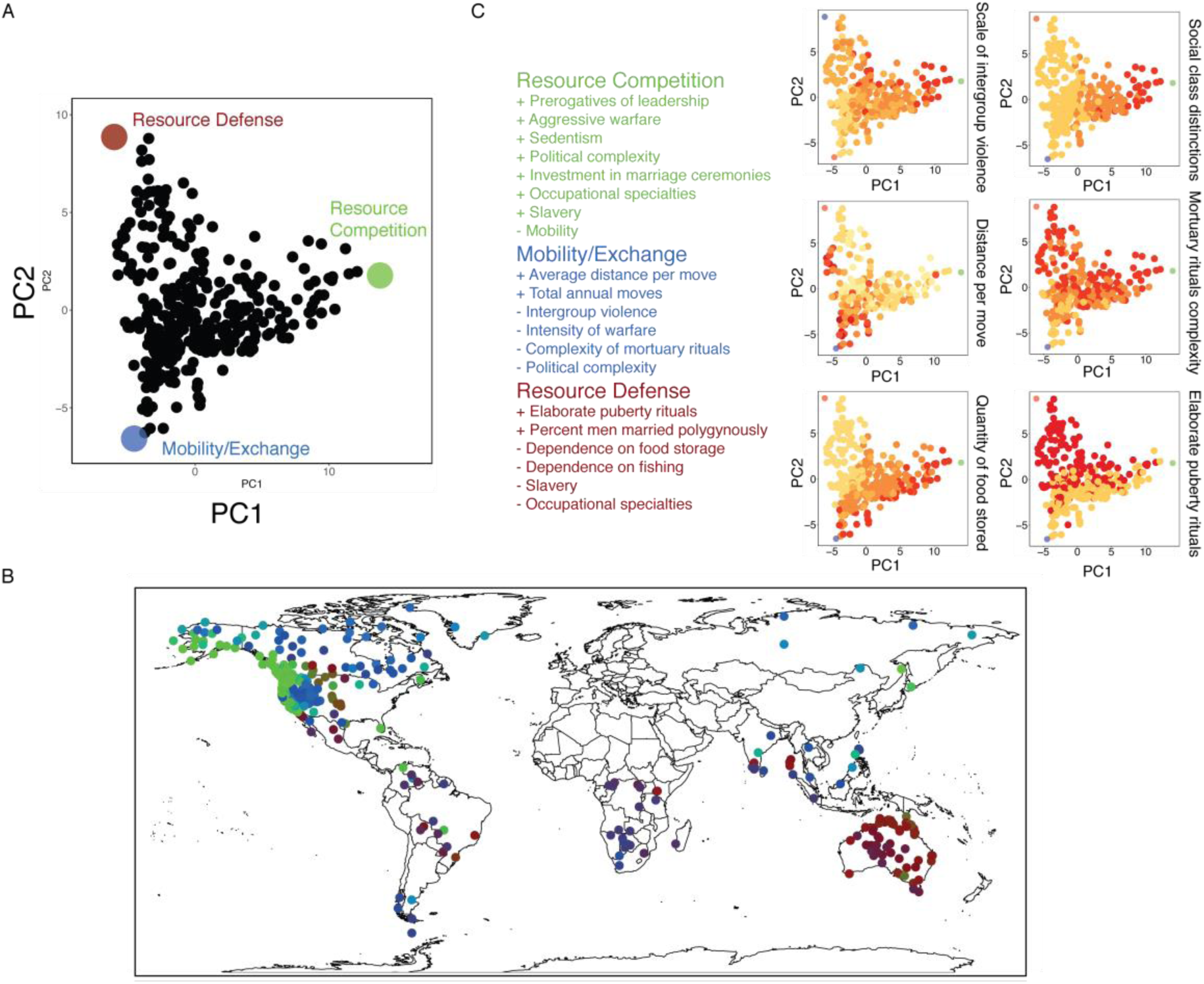
ParTI analysis of *BHG* dataset suggests three tasks. (A) Data plotted on the first two principal components of *BHG* dataset is well-described by a triangle (p<10^-3^). The three vertices of the triangle are hypothesized to be adaptive combinations of traits for three tasks. (B) Cultures are color coded according to their distance from each vertex on a geographical map. (C) List of traits associated with cultures near each vertex. Sign (+/-) indicates positive or negative enrichment. Six of the traits are displayed on the data distribution, with red (yellow) corresponding to a high (low) level of the trait. For example, *social class distinctions* are strongest close to the resource-competition vertex, whereas *elaborate puberty rites* occur predominantly near the resource-defense vertex.

### Characterization of vertices for *BHG*

The hunter-gatherer cultures closest to the Resource-Competition vertex are the stratified societies of the North-American Pacific Northwest, with strong enrichments for *prerogatives of leadership, intense and aggressive warfare* and *political complexity*, as well as *sedentism* (Figure 3C). As with the Austronesian cultures, the hunter-gatherers that are the closest to the Mobility/Exchange vertex come from areas with the lowest resource density – the arctic tundra and deserts. This vertex is enriched with *high mobility, low inter-group violence* and *low religious and political complexity*. Last, the cultures closest to the Resource Defense vertex are Australian cultures from the relatively fertile Arnhem Land. Closeness to this vertex is associated with *low dependence on food storage, elaborate male puberty rituals*, and other traits that relate to *marriage patterns*.

We next asked whether the tasks of the three vertices found in the two datasets might be similar. This comparison is challenging due to the small overlap in traits between the two datasets, and the differences in subsistence strategies. We note that in both datasets, the Resource Competition vertex is enriched with traits that relate to territorial conquest and social dominance and the cultures near the Mobility/Exchange vertex live in areas with lower resource density. We tentatively conclude that the vertices might correspond to comparable tasks, and used information from both datasets to provide an interpretation of these tasks.

### Economic defendability theory and the tasks

After defining the three tasks, we sought a theoretical framework that can unite their understanding. We find that the three tasks can be interpreted using economic defendability theory, a classic theory from human behavioral ecology which relates variation in territorial behavior with differences in resource density and predictability (29). This theory observes that only when a resource is dense and predictable is it worthwhile to engage in costly behaviors to defend it. Otherwise, other behaviors, such as mobility and exchange, will be preferred. Economic defendability was first used to explain the evolution of territorial systems in birds (30), and has been applied to study-small scale human societies (29,31,32).

In agreement with economic defendability theory, the Mobility/Exchange vertex, which is associated with cultures that live in areas with low resource density, is enriched with lack of competitive cultural traits. The cultures nearest this vertex are highly mobile; their reliance on trade or wage labor may compensate for poor and unpredictable resources; prevalence of piercing may be explained by findings that extreme rituals promote pro-sociality and inclusive social identity (33); and their large ritual groups may reflect the need to reinforce cultural identity in the face of dispersal (34). For these reasons, we conclude that this vertex includes traits that are adaptive for the task of mobility and exchange.

Conversely, when resources are sufficiently dense and predictable, territorial strategies should emerge. Territorial strategies require two separate tasks (35) – resource defense and resource competition. The Resource Competition vertex is enriched in both datasets with socially competitive traits, which are associated with conquest and colonization, whereas the Resource Defense vertex is associated with traits that are more cooperative.

Resource competition may become more important than resource defense when there are strong resource gradients (36), unpredictable natural disasters (37) or due to population pressure (38). The enrichments for hierarchy-related traits near the Resource Competition vertex may be due to the advantages of hierarchy for group-coordination (39) or for outgroup exploitation (40).

As an example, the difference between the tasks can be illustrated by the headhunting practices in two phylogenetically close cultures that lie on two sides of the same island: *Laboya* from the western part of the island of Sumba, and *East Sumba* from its less fertile eastern part. *Laboya* lies close to the Resource Defense vertex while *East Sumba* lies close to the Resource Competition vertex. Both cultures used to practice headhunting (the practice of killing people to obtain their heads); however, in West Sumba the motivation for headhunting was reported to be defense of traditional territories, whereas in East Sumba its motivation was reported to be territorial expansion (41), in accordance with the proposed tasks for these vertices.

## Discussion

In this study, we tested the hypothesis that functional tradeoffs are important for adaptation at the level of societies, by using an approach from systems biology to infer tasks and tradeoffs and applying it multivariate datasets of cultural traits. We find a triangular distribution in trait space in two independent cultural datasets, for Austronesian cultures and modern hunter-gatherers. This triangle suggests a fundamental tradeoff between three tasks. Near each vertex are geographically and linguistically distinct cultures that have shared features. These features provide clues about the tasks. Using theory from human behavioral ecology, we identify the three tasks as mobility/exchange, resource defense and resource competition. We detect specific combinations of cultural traits that are predicted to be adaptive for each task. The present approach can in principle be used to infer tasks and tradeoffs for other cultures using multi-variate datasets.

The results provide numerous hypotheses regarding the adaptive role of cultural traits with respect to these three putative tasks. While the functional role of many of the traits fits well with the theory of economic defendability, other suggested associations are surprising. For example, the association of *taboo* with resource competition rather than resource defense suggests further investigation.

An important caveat is that by itself, ParTI cannot test whether a specific trait provides adaptive value, or, instead, whether the trait is linked with another trait that provides adaptive value. These hypotheses should therefore be tested in future studies that employ methods such as phylogenetic analysis or experimental psychology. Such studies can, in turn, help refine our understanding of the tasks and trade-offs that shape the cultural evolutionary process.

The ParTI approach does not depend on the evolutionary mechanisms at play, nor does it suppose that a culture is perfectly adapted to its environment. It provides a quantifiable, geometrical signature for generating hypotheses on the adaptive tasks of human social organization and cultural evolution. The empty space around the triangle that describes the datasets corresponds to combinations of cultural traits that could *a-priori* have existed, but are not found in the data. For example, a combination of small ritual groups (as in resource defense vertex) together with reliance on trade/wage labor (as in mobility/exchange) is rare in the *Pulotu* dataset. ParTI, adjusted for phylogeny and spatial diffusion, explains both empty space and co-occurring traits based on tradeoffs between adaptive tasks.

ParTI might be useful to complement other methods of cultural evolutionary analysis. It overcomes some of the difficulties of existing approaches. One difficulty is in the interpretation of correlations between a given cultural trait and environmental factors (42). For example, collectivist cultural values are correlated with pathogen prevalence, which led to the hypothesis they may represent an anti-pathogen defense strategy (43). Interpreting such environmental associations is, however, often ambiguous, since they often relate to the latitudinal gradient and may be explained by other factors (12,42). ParTI uses multivariate data, rather than single traits, allowing stronger statistical inference and the ability to adjust for confounding geographical and phylogenetic factors.

ParTI might also address some of the limitations of other approaches to model the function of cultural traits, namely optimality models from behavioral ecology (44,45) and co-evolutionary analyses (46,47). Optimality models require information on costs and benefits that is not always available. Co-evolutionary models follow traits along phylogenetic trees, constructed for example from languages. They have been used to study associations such as the link between human sacrifice and social stratification (23) and the link between livestock and descent (48). Such models, however, require strong phylogenetic signals and are not applicable to traits that have weak phylogenetic transmission.

The ParTI approach can apply to any cross-cultural dataset, and thus be used to test hypotheses on the tradeoffs in cultural adaptations. It provides a top-down way to provide hypotheses on the function of cultural traits and the trade-offs that shape cultural variation.

## Author Contributions

OK and UA conceived and performed the research. OK and UA wrote the manuscript.

## Additional Information

Competing Interests: The authors declare no competing interests.

