## appendix for "Tradeoffs and cultural diversity"

### Table of Contents

**SUPPLEMENTARY INFORMATION – TRADEOFFS AND CULTURAL DIVERSITY ... 1**

Supplementary Information ... 2

*Supplementary Information 1. ParTI theory and applications. ... 2*
*Supplementary Information 2. Preparation of Pulotu cultural trait data. ... 4*
*Supplementary Information 3. Preparation of Binford's Hunter-Gatherer (BHG) cultural trait data. ... 7*
*Supplementary Information 4. Number of significant principal components. ... 9*
*Supplementary Information 5. Statistical significance of triangles. ... 11*
*Supplementary Information 6. Trait regression according to distance from vertices. ... 14*
*Supplementary Information 7. The relation between island type and distance from archetypes for Polynesian and Micronesian cultures. ... 17*
*Supplementary Information 8. ParTI analysis of Schwartz cultural value orientations. ... 18*
*Tree1. Language tree for Pulotu (informative branch lengths) ... 20*
*Tree2. Language tree for Pulotu (uninformative branch lengths) ... 21*
*Tree3. Language tree for BHG (uninformative branch lengths) ... 22*
*Table S1. Vertices for Pulotu ... 23*
*Table S2. Vertices for BHG ... 24*
*Table S3. Features used from Pulotu ... 25*
*Table S4. Features used from BHG ... 25*
*Table S5. Limit of data. ... 25*
*Table S6. Traits associated with vertices for Pulotu ... 25*
*Table S7. Traits associated with vertices for BHG ... 25*
*Table S8. Distances from archetypes for Pulotu ... 25*
*Table S9. Distances from archetypes for BHG ... 25*
*Table S10. Data for the relation between island type and distance from archetypes for Polynesian and Micronesian cultures. ... 26*

References ... 27

### Supplementary Information

#### Supplementary Information 1. ParTI theory and applications.

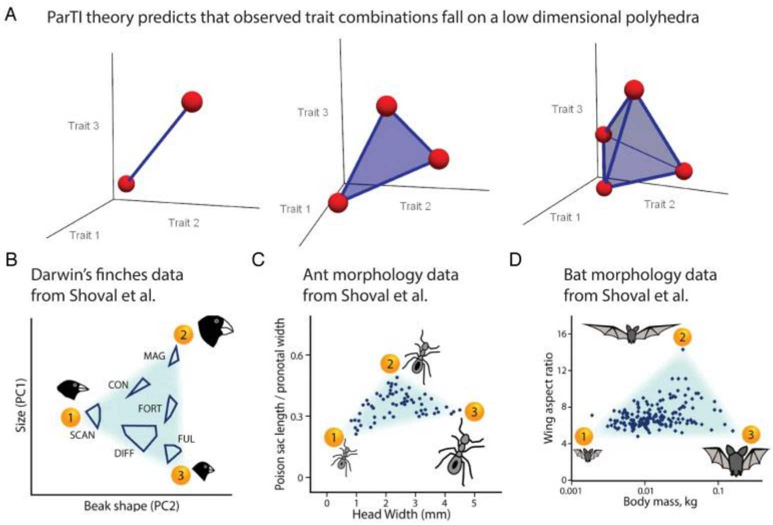


**Figure S1.**

Consider a vector of traits $\vec{T}$ so that fitness results from performance in $k$ tasks, so that performance is given by $k$ different performance functions: $F(\vec{T}) = F(P_1(\vec{T}), P_2(\vec{T}), \ldots, P_k(\vec{T}))$. It is possible that no single trait vector $\vec{T}$ can maximize performance of all performance functions at once, leading to a fundamental tradeoff.

ParTI theory shows that the best trade-off trait combinations fall on low-dimensional shapes in trait space with vertices and flat sides, known as polytopes (such as lines, triangles, tetrahedra and so on). The vertices of the polytope represent the trait combinations that are optimal for each task, that is, vertex $i$ is the trait vector $\vec{T_i}$ that optimizes $P_i(\vec{T})$. For two tasks, the data should fall on a line segment with the two vertices types at the ends; for three tasks, data should fall on a triangle with the three vertices at the corners and so on. Specialists at a task lie near the vertex, and generalists lie in the center of the polytope (Figure S1A). Under certain conditions, when performance at tasks depends in a complex way on the traits, slightly warped polytopes are predicted (*3*). Notably, not all possible traits need to be measured- only enough traits to reliably define the polytopes (*1*).

The main use of ParTI is to infer the number and nature of tasks that played a role in the evolution of the system. ParTI was implemented in an algorithm (*2*) that tests: (i) whether the data is low dimensional, (ii) whether in the low dimensional space, the data is well described by a polytope, (iii) which traits of the data are found in the data points closest to each vertex- to provide clues for the task. Statistical tests assign p-values to each of these steps.

Shoval et al. showed that the classic measurement of bird morphology of Darwin's finches (*4*) by the Grants fall on a low dimensional triangle (*1*) (Figure S1B). This triangle is defined by three vertices that correspond to specialists in eating insects/nectar, large/hard seeds and small/soft seeds. Other examples discussed in the Shoval et al. manuscript include measurements of leaf cutter ant morphology (Figure S1C) and bat wing morphology (Figure S1D), as well as bacterial gene expression.

Subsequent work used ParTI theory to study various biological datasets, including cancer gene expression (*2*), life history traits (*5*), ammonite fossil morphology (*6*) single cell gene expression (*7*) and other datasets (*8, 9*).

The same reasoning explains why ParTI theory should apply to adaptive cultural variation. The adaptive importance of human culture, as well as processes that may select for adaptive cultural traits, have been extensively studied (*10–24*). A cultural trait is adaptive if it increases performance in a task that is relevant for fitness. For example, cultural traits that are relevant for social organization can cause groups to be more successful than others in inter-group competition (*11*, *25*, *26*) or serve as anti-pathogen defense strategies (*27*).

### Supplementary Information 2. Preparation of *Pulotu* cultural trait data.

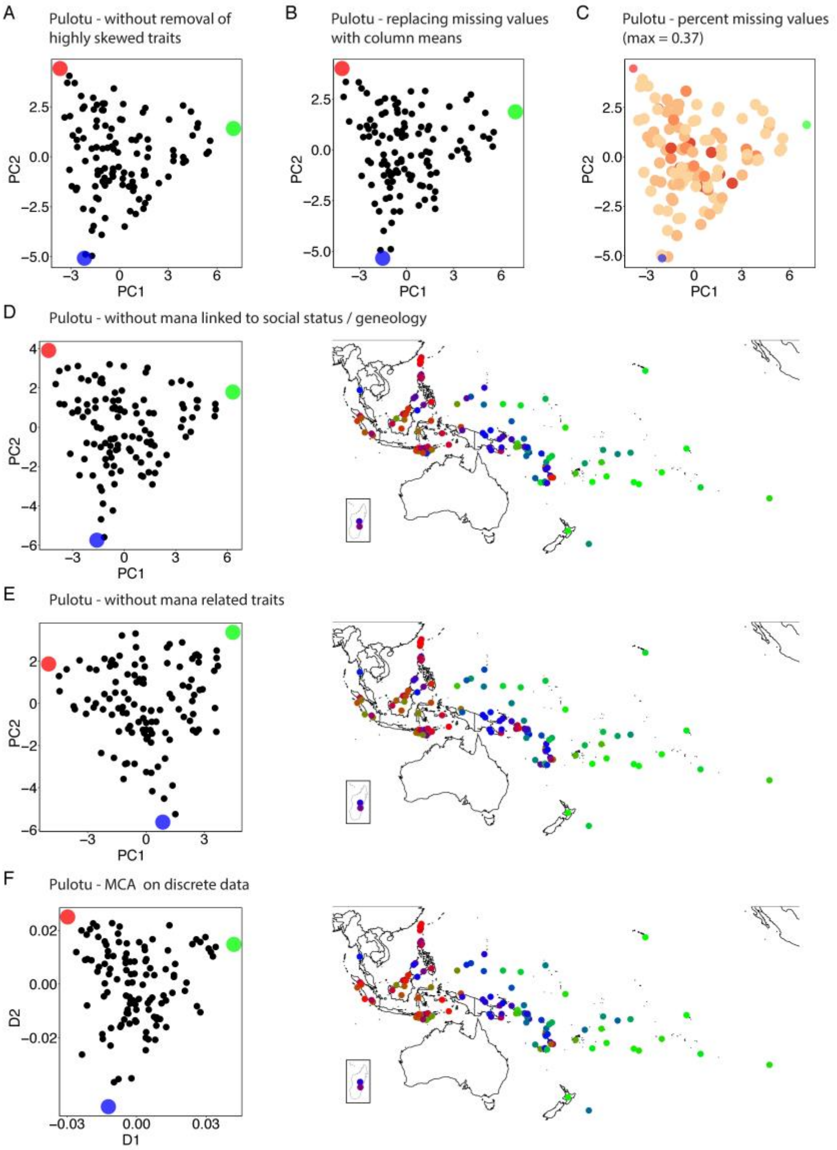


**Figure S2.**

*Trait selection. Pulotu* (*28*) was retrieved from the website: https://pulotu.econ.mpg.de/. It contains traits for 116 Austronesian cultures. Among these traits we used 44 traits that are cultural (i.e. not physical) and belong to the traditional time focus as defined by Watts et al. (*28*), which we supplemented with the human sacrifice and social stratification traits from Watts et al. (*29*).

Because highly skewed traits may create outliers in the PCA, we discarded three traits whose skewness is larger than 4 - *Forces of nature are controlled by or imbued with the supernatural* (v35), *Supernatural punishment for impiety* (v44) and *Kinship Tapu* (v52). These features are unimportant in the first two principal components and when we include them, we get a very similar triangle (Figure S2A). We also removed from the data a single outlier culture (Eastern Toraja). We accounted for the removal of this single outlier culture in the calculation of the significance of the triangle. The traits that are used in the ParTI analysis are available in (Table S3).

*Missing values.* The resulting dataset contains ~13% missing values, which were imputed by taking the mean of 11 multiple imputations with predictive mean matching. The imputation was calculated using the r package mice. The triangle is not an artifact of the imputation, as a similar triangle results from simply replacing missing values with column means (Figure S2B).

For cultures that have a high percentage of missing values, there is some uncertainty as to where they should fall on the triangle. Therefore we removed two cultures with a very high (>70%) percentage of missing values - *Kosrae* and *Goodenough Island*. These two cultures fall in the center of the triangle and therefore do not influence the observed geometry. This leaves the maximal percent of missing values for a culture to be 37%. Importantly, the cultures with a high percentage of missing values do not define the triangle, that is, most of the archetypical cultures do not have a high percent of missing values (Figure S2C).

*Robustness to mana related traits.* The most significant enrichments are traits that represent various aspects of mana - mana related to social influence or technical skill, mana as a spiritual or religious concept and mana as a personal quality (v56-v58). The traits include also two sub-traits of mana as a personal quality - mana and social status and mana linked to genealogy (v59-v60). The first three traits (v56-v58) are the three most important contributors to the first principal component, while the last two contribute to the second principal component. Since the traits (v59-v60) are defined only when the trait (v58) equals one (v58=1), it is not straightforward to impute them with a meaningful value when (v58=0). We therefore asked to what extent excluding these traits affects the observed geometry. We found that when we exclude these traits we get a triangle that is similar to the original triangle, with similar distances from the vertices for different cultures - distances from vertices have a correlation larger than 0.97 for each of the three distance vectors (Figure S2D). Because of this, and since we think that these traits are functionally significant (see main text), we chose to keep them in the analysis. Other than this, any combination of *Pulotu* traits should be plausible a-priori.

We then further asked how robust the observed triangle is with respect to *all* mana related traits. To test this, we removed all of the five mana related traits from the data and performed the ParTI analysis. We find that although the first two principal components are slightly different, we get a triangle that is similar to the original triangle (Figure S2E), with similar distances from the vertices for different cultures - distances from vertices have a correlation larger than 0.9 for each of the three distance vectors. This shows that the triangle is robust with respect to removal of the mana related traits.

*Multiple correspondence analysis.* We tested whether the observed geometry is sensitive to the method of dimensionality reduction. Because the *Pulotu* dataset is discrete, we used *multiple correspondence analysis (MCA)* and then performed the ParTI analysis on the first two output components (MCA was performed using the r package homals). The imputed data was prepared for the MCA by taking the median of the 11 multiple imputations (instead of the mean). We find that the geometry is explained by a triangle that is similar to the original triangle (Figure S2F), with similar distances from the vertices for different cultures - distances from vertices have a correlation larger than 0.95 for each of the three distance vectors.

### Supplementary Information 3. Preparation of *Binford's Hunter-Gatherer* (BHG) cultural trait data.

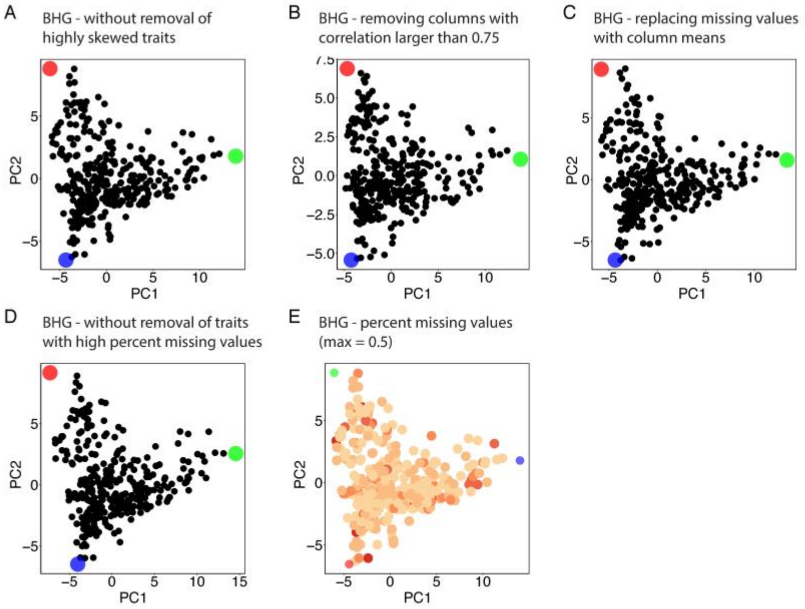


**Figure S3.**

*Trait selection. Binford's Hunter-Gatherer* (*30*) dataset *LRB.csv* was retrieved from http://ajohnson.sites.truman.edu/data-and-program/ (data is also available in the r package binford). It contains 200 traits for 339 hunter-gatherer societies. From these traits, we extracted 88 cultural traits that are numeric and ordinal. These traits are a mixture of discrete and continuous traits. To guarantee that the resulting geometry is not influenced by variables that must sum up to a constant, we removed the columns for percent reliance on hunting/gathering. This dataset contains several highly correlated traits (Pearson or Spearman correlation > 0.95) which we removed, with skewed traits preferentially removed. As with the *Pulotu* dataset, after this procedure we also removed highly skewed traits (skewness > 4). There is only one such trait - *hunt*, which codes for the degree to which males conduct hunting, relative to females. When we include this trait, we get a very similar triangle (Figure S3A). The triangle also does not strongly depend on the threshold for the removal of the correlated traits – for example, removing traits with spearman or Pearson correlation larger than 0.75 yields a similar triangle, with correlation with distances from the vertices larger than 0.86 (Figure S3B). After this preprocessing, 66 traits remain (Table S4).

*Missing values.* The remaining traits have 20% missing values. To reduce the percent of missing values, we removed traits that have more than 50% missing values (10 traits). Removing these traits has a very small effect on the triangle (Figure S3C), and it reduces the percent of missing values to 12%. We imputed these missing by using the same method that was used for the *Pulotu* dataset. As with the *Pulotu* dataset, here as well the triangle is not an artifact of the imputation and a similar triangle results from replacing missing values with column means (Figure S3D), and the cultures with a high percentage of missing values do not define the triangle (Figure S3E).

### Supplementary Information 4. Number of significant principal components.

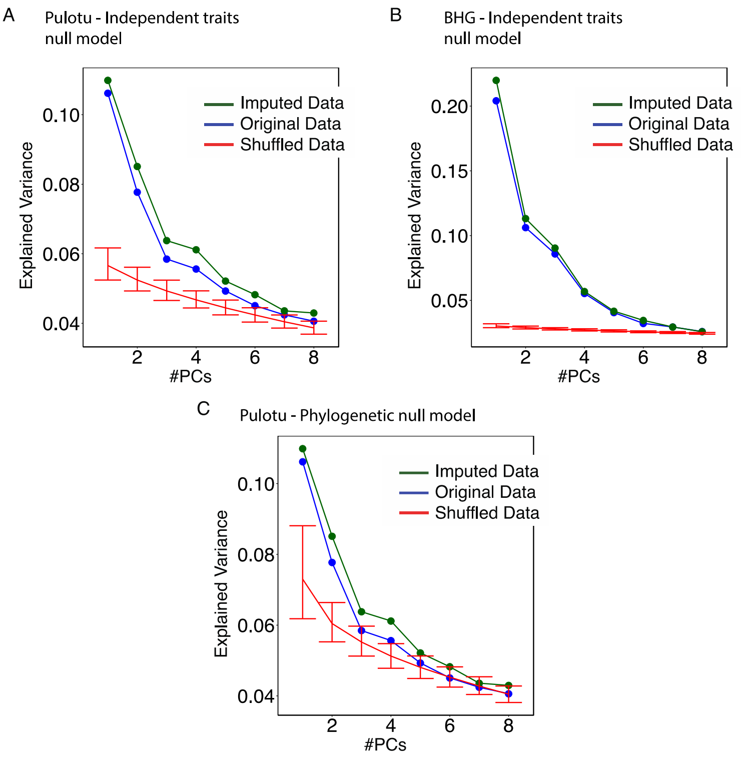


**Figure S4.**

We inferred the effective dimensionality of the datasets – the number of principal components that cannot be attributed to random fluctuations. For this, we compare the variance that is explained by each principal component to the variance that is explained by a random null model of the data.

A basic null model assumes that each trait is completely independent from other traits. Under this null model, random datasets can be generated by simply shuffling the columns of the original dataset. We call this null model the *independent traits null model*. We used the *independent traits null model* to generate 1000 random datasets. For each of these datasets we performed principal component analysis and then calculated means and 95% confidence intervals for each principal component (Figure S4AB). The *independent traits null model* reveals high non-random dimensionality in *Pulotu* and in *BHG*.

However, this null model does not account for the possible dependencies in the data that may result from random processes, such as phylogeny or spatial diffusion. These random processes may generate correlations between traits that are independent from each other.

To infer the non-phylogenetic dimensionality of the data, we generated a *phylogenetic null model* where traits are acquired and lost at random. This null model requires phylogenetic information on culture diversification with informative branch lengths, which is available for 93 cultures from the *Pulotu* dataset[26]. We therefore constructed this null model only for the *Pulotu* dataset.

We constructed the *phylogenetic null model* as follows. Because all the traits in the *Pulotu* dataset are discrete, we used a discrete trait evolution model where each trait can get values $p_1,\ldots p_n$ and there is a transition probability between every two traits $p_i \rightarrow p_j$. These transition probabilities define a transition matrix *P*. We fit each transition matrix from the data by maximal likelihood discrete trait reconstruction (using the ace function in the r package ape). With the transition matrix, we simulate the trait values using the function sim.history in the r package phytools. Using this method, we can generate datasets with random independent traits that maintain the phylogenetic structure of the *Pulotu* dataset.

We used the *phylogenetic null model* to generate 1000 random datasets. For each of these datasets we performed principal component analysis and then calculated means and 95% confidence intervals for each principal component (Figure S4C). This analysis reveals that only the first two principal components of the *Pulotu* dataset are significant beyond phylogenetic fluctuations. For this reason, we chose to use only the first two principal components of each dataset for the analysis.

This analysis suggests that while the first two principal components only explain a small percent (~20%) of the variance in the *Pulotu* dataset, the rest of the variance in the dataset may be attributed to noise or random phylogenetic effects. By increasing the number of features measured, however, it may be possible to discover more tasks and tradeoffs in this dataset.

Since a phylogenetic tree with informative branch lengths does not exist for the *BHG* dataset, we could not apply this approach to test its dimensionality. An inspection of the traits that are enriched in each of the principal components of the *BHG* dataset showed that the trait with the highest score in the third PC is the *existence of mounted hunters*. Moreover, the societies were bimodally distributed on this PC, with a distinct cluster of mounted hunters. This indicates that the third PC separates cultures according to whether they had access to mounted horses, and is therefore not relevant for our analysis. For this reason, and to compare the results to the analysis of the *Pulotu* dataset, we used only the first two components to analyze the *BHG* dataset.

### Supplementary Information 5. Statistical significance of triangles.

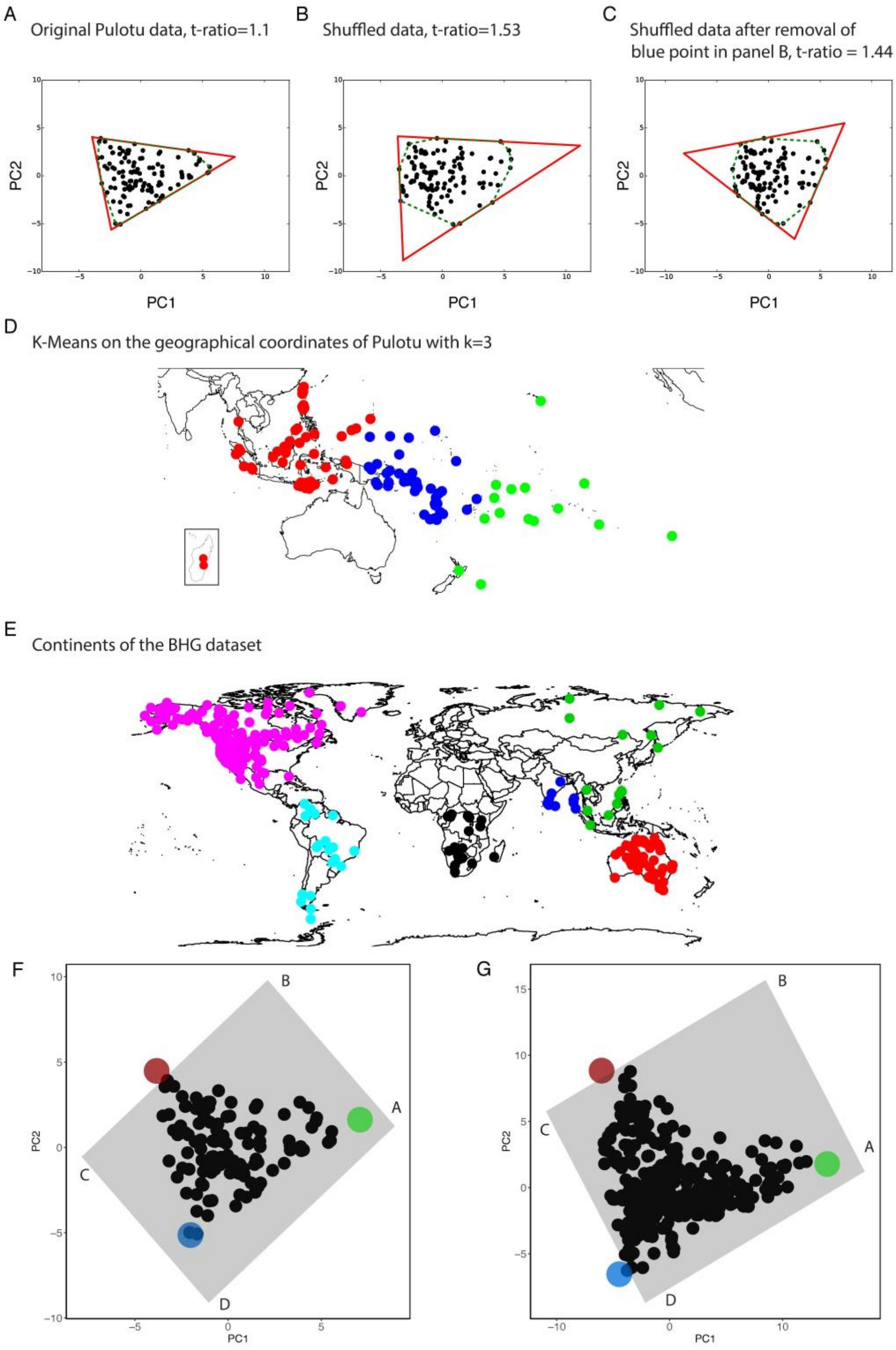


**Figure S5.**

The statistical significance of the observed triangle geometry of each dataset was estimated by using the t-ratio test as specified by Shoval et al. (*1*) (See Figure 3 in that paper for an example) and Hart et al. (*2*). The t-ratio is the ratio between the convex hull of the data and the minimal enclosing triangle, and the t-ratio test computes a p-value by comparing the t-ratio of the data to t-ratios of shuffled data. To account for the removal of an outlier culture in the *Pulotu* dataset we also performed on the *Pulotu* dataset a test that calculates the t-ratio of the shuffled data as the minimal t-ratio after the removal of any observation. We illustrate the calculation of the t-ratio in Figure S5ABC. We performed geometrical calculations in python with the package cv2, with the enclosing triangle calculated by the function minEnclosingTriangle and the convex hull calculated by the function ConvexHull.

The t-ratio of the *Pulotu* dataset is $t_{ratio}=1.1$ and the t-ratio of *BHG* is $t_{ratio}=1.12$. First, we randomized data points by shuffling the observed *x* and *y* coordinates the data points (as specified in Shoval et al. (*1*) and Hart et al. (*2*)), assuming that all points are independent. The observed t-ratios are lower than the minimal t-ratios of 1000 shuffles so the p-value for both triangles is $p<10^{-3}$.

We then tested whether the observed geometry may have been affected by the principal component analysis (PCA). To do so, we randomized the datasets before the PCA by shuffling the columns, and then calculated the t-ratio of the first two principal components. Again, we find that the observed t-ratios are lower than the minimal t-ratios of 1000 shuffles for both datasets ($p<10^{-3}$).

To account for possible effects of phylogeny or spatial diffusion on the observed t-ratios, we performed tests where we shuffle data points only within certain clusters and measure the resulting t-ratio. In the *Pulotu* dataset, for example, there is a clear spatial trend where western cultures tend to be closer to the Resource Competition vertex, eastern cultures tend to be closer to the Resource Defense vertex and central (Melanesian) cultures tend to be closer to the Mobility/Exchange vertex. This may raise the concern that these three clusters affect the observed geometry of the data. We therefore generated shuffled data where *x,y* could only be shuffled between points within the same cluster (clusters were calculated using k-means – see Figure S5D). For 1000 such shuffles we get $p<10^{-3}$ for the *Pulotu* dataset.

Similarly, there are spatial trends in the *BHG* dataset, where most Australian cultures lie near the Resource Defense vertex and most of the cultures near the Resource Competition vertex are from North America. We therefore generated shuffled data where *x,y* could only be shuffled between points within the same continent (Figure S5E). For 1000 such shuffles we get $p<10^{-3}$ for the *BHG* dataset.

We further tested whether the observed t-ratio may result from some other phylogenetic effect. To account for this, we computed the t-ratio of the 1000 randomized datasets generated by the *Pulotu* phylogenetic null model (for details on this null model see *Number of significant principal components*, traits were prepared as in *Preparation of Pulotu trait data*). These datasets only include the cultures for which phylogenetic data is available (93 cultures). We find that 2/1000 of the randomized datasets have a smaller t-ratio than the observed data (p=0.002), and 8/1000 have a smaller t-ratio after removing any single outlier that may give a lower t-ratio (p=0.008). However, these randomized datasets have a low t-ratio because they are sensitive to outliers that are far away from the data, which artificially increases ratio of the convex hull to its enclosing triangle. The t-ratio is therefore highly sensitive to the removal of such outliers. Indeed, when we calculate the t-ratio as the maximal t-ratio after the removal of any observation, we get that the t-ratio of the observed data is again the lowest ($p<10^{-3}$).

To gain insight into the tasks, we characterized the trait-combinations that were missing (the empty trait-space). For this we inferred the projection to PC1, PC2 of all possible trait combinations for *Pulotu* and *BHG* (Figure S5FG, see Tables S5 for the trait combinations that define the corners of the inferred projection). Both datasets occupy a similar space in PC1, PC2, with two trait combinations clearly missing – the corners *B,C.* In *Pulotu,* Corner *B* corresponds to a hypothetical culture with traits that relate to mana and social hierarchy and with low conflict within communities, as well as with high conflict with other communities and cultures, with large ritual groups and with the belief that actions affect afterlife. Corner *C* represents cultural that rely on trade, but also have high levels of conflict between communities and with other cultural and practice headhunting. In *BHG*, Corner *B* is also associated with social hierarchy and with aggressive warfare, but also with initiation rituals, with organized revenge and with the segregation of boys prior to puberty. Corner *C* represents cultures without warfare, but also with polygyny and with initiation rituals. Most importantly, the empty corners represent plausible trait combinations and are therefore not an artifact of the structure of the dataset.

### Supplementary Information 6. Trait regression according to distance from vertices.

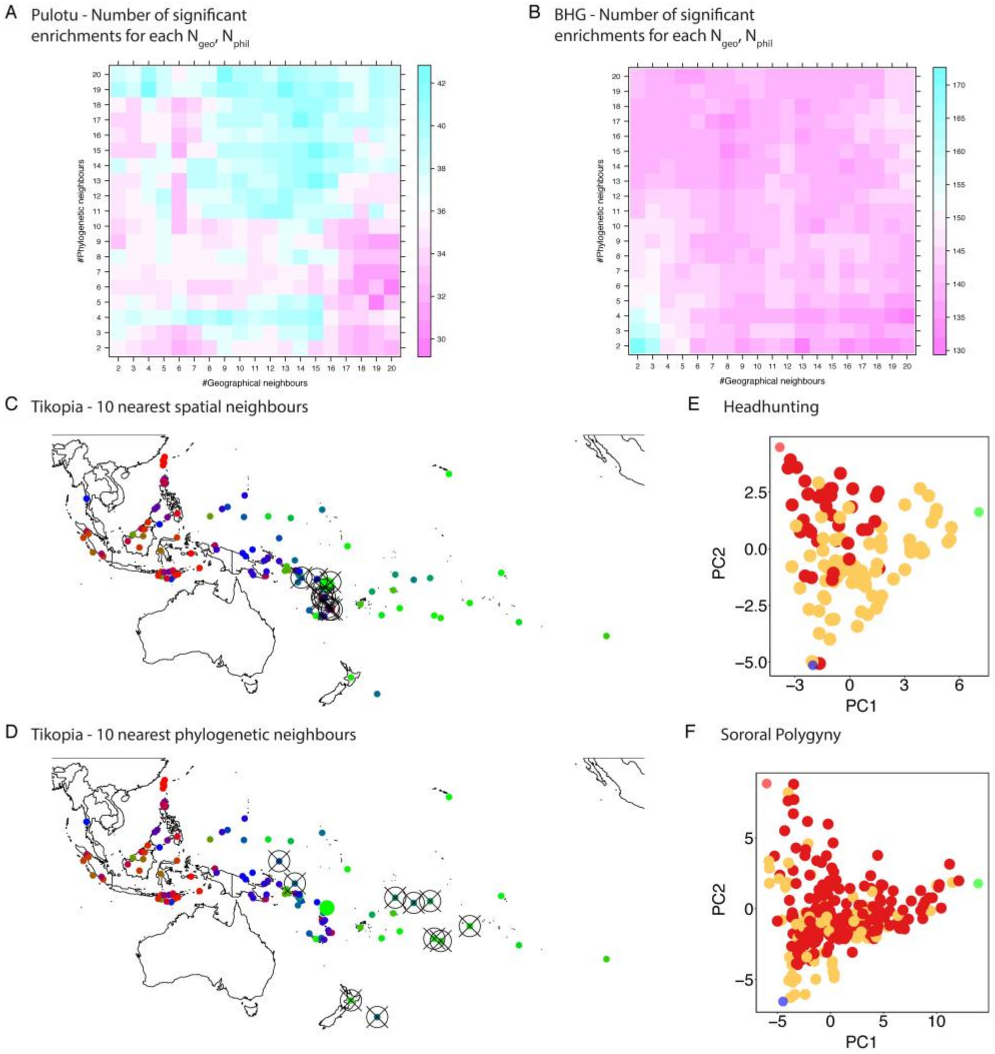


**Figure S6.**

We expect that features that are functional for a task that is represented by a vertex will increase in level or frequency (for binary traits) near that vertex. We denote that such a feature is *enriched* near a vertex. To test for feature enrichment, we ask to what extent does the distance from a vertex, measured as Euclidean distance in the space of PC1, PC2, explain the frequency or level of that feature. We therefore test feature enrichment by performing regression, where the response variable is the feature and the predictor variable is the distance from vertex.

We first generated features from the cultural traits of each dataset. These features include all the original cultural traits of each dataset (before imputation). Traits that have skewness larger than 2 or kurtosis larger than 2 were binarized as either larger/smaller than mean, and traits that have only 3 possible values (such as piercing in *Pulotu*) were binarized as two variables - less than the maximal value or more than the minimal value (marked as LOW and HIGH, respectively). This preparation results in 57 candidate features for *Pulotu* and with 85 candidate features for *BHG*.

It is important to note that a certain feature may be enriched near a vertex just because of random processes such as phylogenetic transmission or spatial diffusion. For example, in the *Pulotu* dataset, the cultures that are closest to the Resource Competition vertex are all Polynesian. Therefore, it may be that some cultural traits will appear more frequently near that vertex just because they relate to the common heritage of Polynesian cultures. To adjust for this phenomenon, we add phylogenetic and geographical covariates to our regression. The geographical covariate for a culture is the average value of the feature for the $N_{geo}$ nearest geographical neighbors, and the phylogenetic covariate is the average value of the feature for the $N_{phyl}$ nearest phylogenetic neighbors. We measure phylogenetic distance as the distance in branch lengths, with ties broken according to geographical distance. This method is similar to the method used by Botero et al. (*31*) to adjust for geographical effects. We set $N_{geo}=N_{phyl}=10$ as in Botero et al. (*31*). The number of significant enrichments for each dataset is not very sensitive to the exact values of $N_{geo}$, $N_{phyl}$ around this value (Figure S6AB).

As an example, consider the culture *Tikopia* in the *Pulotu* dataset. *Tikopia* is a Polynesian outlier culture, that is, despite being nearer in terms of language phylogeny to Polynesian cultures, it is located outside Polynesia and is spatially nearer to Melanesian cultures. Therefore, the set of 10 nearest phylogenetic neighbors is distinct for this culture from the set of 10 nearest spatial neighbors (Figure S6BC).

We thus use the following regression formula for each feature. Let **FEATURE**, **ARCH_DIST, NN_PHYL, NN_SPATIAL** be vectors representing feature values, distance from vertex, and average phylogenetic/spatial values for N-nearest neighbors. The regression is:

**FEATURE ~ -ARCH_DIST + NN_PHYL + NN_SPATIAL**

We then extract a p-value for each coefficient using the t-statistics of the coefficients. We run this regression for each feature and for each of the three vertices and correct the p-values for multiple hypotheses testing using FDR. Using a threshold of 0.02 for the FDR-adjusted p-values, we find in the *Pulotu* dataset 35 enrichments (Table S6). Because of the large number cultures in the *BHG* dataset, we used a lower enrichment threshold of adjusted p-values smaller than 0.0002. Using this threshold, we find 79 enrichments (Table S7).

Adjusting for spatial and phylogenetic effects removes several enrichments that would otherwise be significant. For example, the headhunting trait in *Pulotu* seems strongly enriched near the Resource Defense vertex (Figure S6E); however, its prevalence is better explained by spatial diffusion. Another example is the enrichment for sororal polygyny in *BHG*, which is best explained by phylogeny (Figure S6F).

This approach can be applied to study functional hypothesis on new traits. To measure whether a cultural trait has an adaptive role that is relevant for the tasks identified in this study, it is necessary to measure it in the relevant cultures (either *Pulotu* or *BHG*). Then it is possible to test whether it may be adaptive for one of the tasks by applying the regression formula using phylogenetic/spatial information and the distances from the vertices of each culture, which is available in (Table S8, S9).

Last, the PCHA algorithm used for inferring the simplex (see methods) outputs the vertices as points in higher dimensional space. These points (rounded) are attached in (Table S1, S2), and can be interpreted by using the codebooks of *Pulotu* and *BHG* (note that some of the variables are reverse-coded).

### Supplementary Information 7. The relation between island type and distance from archetypes for Polynesian and Micronesian cultures.

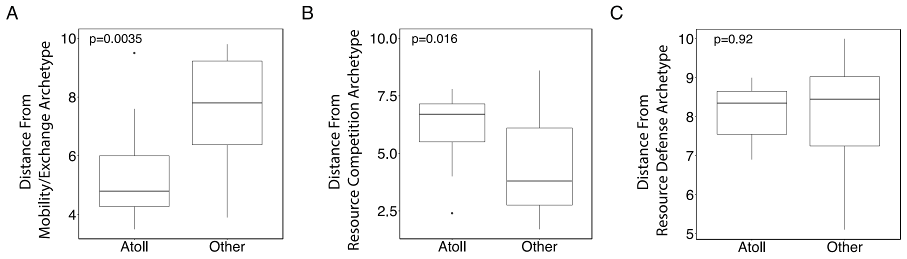


**Figure S7.**

The relation between island type and social organization among Pacific cultures has been long noted by anthropological studies, such as by the classic studies of Sahlins (*32*) and Goldman (*33*). Sahlins noted that in low atolls, where resources are scarce and unpredictable, social organization is the least stratified (*32*). We hypothesized that atoll societies will be closer to the Mobility/Exchange archetype. To test this, we compared the distances from vertices for 32 Polynesian and Micronesian societies (12 atoll societies and 20 societies that reside on high islands, Figure S7 and Table S10). Those that reside on atolls are (A) closer to the Mobility/Exchange vertex (p=0.0035), and (B) further from the resource competition vertex (p=0.016). There is no significant difference for the distance from the Resource Defense vertex (C).

### Supplementary Information 8. ParTI analysis of Schwartz cultural value orientations.

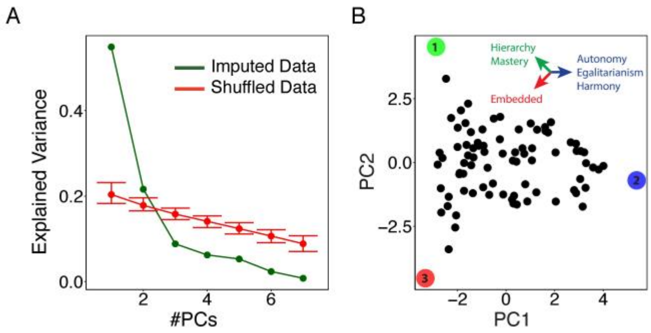


**Figure S8.**

In this study, we detected three basic tasks with tradeoffs that underlie diversity in cultural traits – Resource Defense, Resource Competition and Mobility/Exchange, which we discovered by analyzing cultural traits from Austronesian cultures and from modern hunter-gatherers. A question that arises from the study is how cultural traits may increase adaptive value for these tasks. One way this can come about is by the interaction of culture with individual values – cultural institutions may coordinate individual values across society, and therefore influences individual motivational goals (*34*). We therefore hypothesize that cross-cultural variation in cultural values will also be triangular, with vertices that correspond to the cross-cultural tasks.

To test this, we apply ParTI to a dataset of Schwartz cultural value orientations (*34*) across 80 countries. The data was retrieved from here: https://www.researchgate.net/publication/304715744_The_7_Schwartz_cultural_value_orientation_scores_for_80_countries

It includes the following 7 Schwartz value orientations:

- **Harmony** - fitting into the world as it is
- **Embeddedness** - people are viewed as entities embedded in the collectivity
- **Hierarchy** - hierarchical systems of ascribed roles
- **Mastery** - active self-assertion in order to master, direct, and change the natural and social environment to attain group or personal goals
- **Affective Autonomy** - encourages individuals to pursue affectively positive experience for themselves
- **Intellectual Autonomy** - encourages individuals to pursue their own ideas and intellectual directions independently

- **Egalitarianism -** seeks to induce people to recognize one another as moral equals who share basic interests as human beings

We find that the data is 2 dimensional (Figure S8A) and well described by a triangle (t-ratio = 1.15, p=0.002, Figure S8B). To characterize the vertices, we calculate the pearson correlation between the level of each value orientation and the (reversed) distance from each vertex. We obtain the following correlations:

| Vertex Number | Harmony | Embeddedness | Hierarchy | Mastery | Affective Autonomy | Intellectual Autonomy | Egalitarianism |
|---|---|---|---|---|---|---|---|
| 1 | -0.75 | 0.37 | 0.79 | 0.77 | -0.18 | -0.49 | -0.62 |
| 2 | 0.7 | -0.88 | -0.78 | -0.27 | 0.74 | 0.9 | 0.71 |
| 3 | -0.24 | 0.91 | 0.31 | -0.4 | -0.89 | -0.81 | -0.45 |

Thus we find that vertex (1) is characterized by high Hierarchy, Mastery, vertex (2) is characterized by high Harmony, Affective/Intellectual Autonomy and Egalitarianism, and vertex (3) is characterized by high Embeddedness. We hypothesize that these value orientations may correspond to the resource competition, mobility/exchange and resource defense tasks, respectively.

### Tree1. Language tree for *Pulotu* (informative branch lengths)

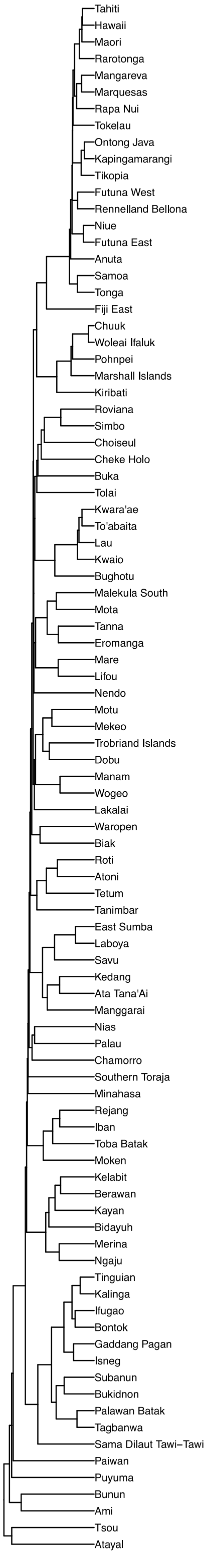

### Tree2. Language tree for *Pulotu* (uninformative branch lengths)

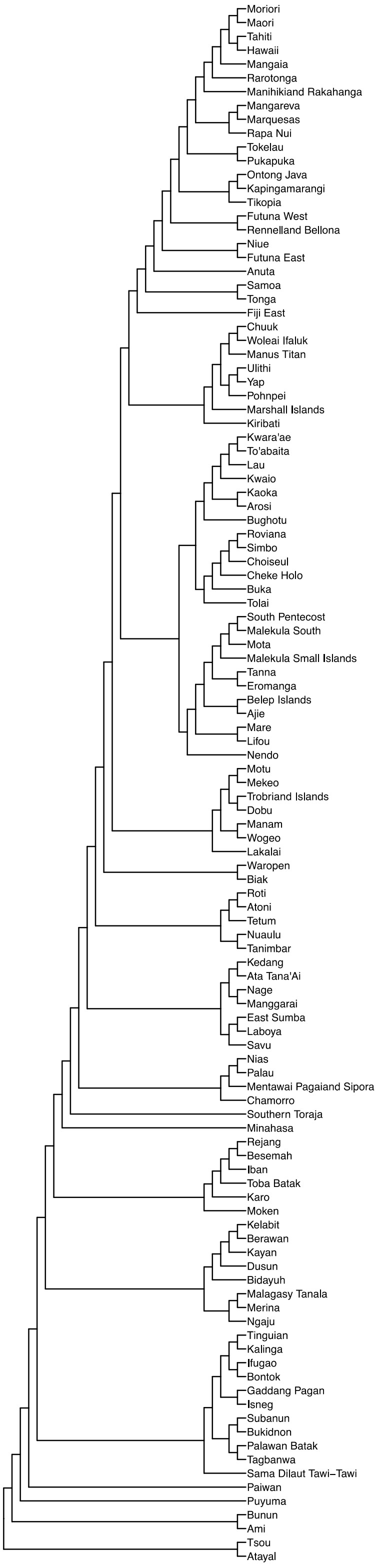

### Tree3. Language tree for *BHG* (uninformative branch lengths)

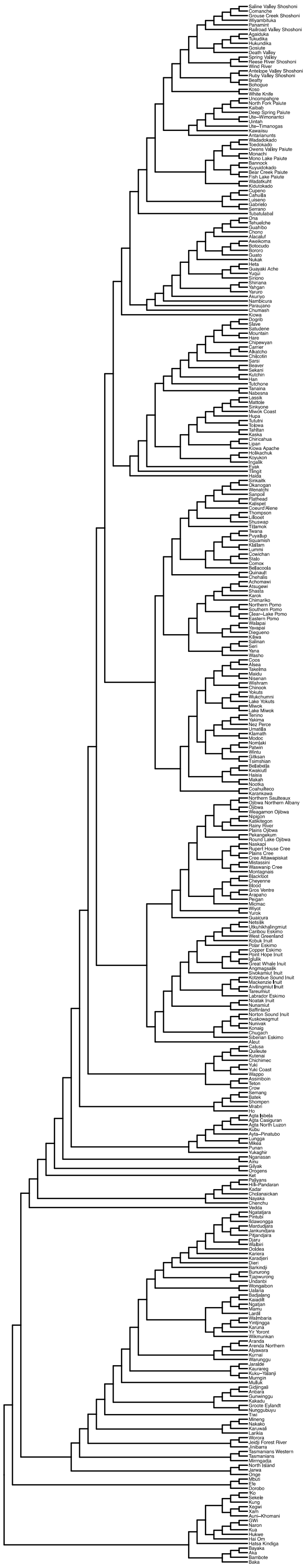

### Table S1. Vertices for *Pulotu*

| Feature | Mobility/Exchange | Resource Competition | Resource Defense |
|---|---|---|---|
| **v14.Conflict_within_communities (reverse coded)** | 3 | 2 | 4 |
| **v15.Conflict_between_communities_of_the_culture (reverse coded)** | 3 | 2 | 1 |
| **v16.Conflict_with_other_cultures (reverse coded)** | 3 | 4 | 2 |
| **v17.Contact_with_other_cultures (reverse coded)** | 1 | 2 | 1 |
| **v24.Agriculture_._Horticulture** | 3 | 4 | 4 |
| **v25.Land.based_gathering** | 2 | 2 | 2 |
| **v26.Animal_husbandry_as_a_source_of_food** | 1 | 1 | 2 |
| **v27.Land.based_hunting_performed_by_individuals** | 1 | 1 | 2 |
| **v28.Land.based_hunting_performed_by_one_or_more_groups** | 0 | 1 | 3 |
| **v29.Water.based_gathering** | 2 | 3 | 0 |
| **v30.Polygamy** | 3 | 1 | 3 |
| **v31.Fishing_and_water.based_hunting_performed_by_one_or_more_groups** | 2 | 3 | 1 |
| **v32.Trade_._wage_labour_as_a_source_of_food** | 3 | 1 | 1 |
| **v37.Nature_Spirits** | 1 | 1 | 2 |
| **v38.Nature_god.s.** | 0 | 2 | 2 |
| **v39.Ancestral_spirits** | 3 | 1 | 2 |
| **v40.Deified_ancestor.s.** | 1 | 3 | 1 |
| **v41.Culture_hero.es.** | 1 | 2 | 1 |
| **v42.God.s.** | 0 | 2 | 2 |
| **v46.One.s_actions_while_living_can_affect_the_nature_of_one.s_afterlife** | 0 | 1 | 1 |
| **v47.The_actions_of_others_after_one_has_died_can_affect_the_nature_of_one.s_afterlife** | 1 | 1 | 1 |
| **v48.Myth_of_man.s_creation** | 0 | 1 | 1 |
| **v49.Primordial_pair** | 0 | 1 | 1 |
| **v51.Social_hierarchy_tapu** | 1 | 1 | 0 |
| **v53.Resource_management_tapu** | 1 | 1 | 0 |
| **v56.Mana_related_to_social_influence_or_technical_skill_** | 0 | 1 | 0 |
| **v57.Mana_as_a_spiritual_or_religious_concept_** | 0 | 1 | 0 |
| **v58.Mana_as_a_personal_quality** | 0 | 1 | 0 |
| **v59.Mana_and_social_status** | 1 | 1 | 1 |
| **v60.Mana_linked_to_genealogy** | 0 | 3 | 2 |
| **v61.Political_and_religious_differentiation** | 2 | 2 | 2 |
| **v63.Headhunting** | 0 | 0 | 1 |
| **v64.Costly_sacrifices_and_offerings** | 0 | 1 | 1 |
| **v65.Size_of_largest_ritual_social_group** | 4 | 3 | 3 |
| **v66.Tattooing** | 0 | 2 | 1 |
| **v67.Scarification** | 1 | 0 | 0 |
| **v68.Piercing** | 2 | 1 | 0 |
| **v69.Genital_cutting** | 0 | 2 | 1 |
| **v70.Tooth_pulling** | 0 | 0 | 1 |
| **v105.Importance_of_Patrilateral_descent** | 2 | 3 | 2 |
| **v106.Importance_of_Matrilateral_descent** | 2 | 2 | 2 |
| **Stratification** | 1 | 3 | 2 |
| **Sacrifice** | 0 | 1 | 0 |

### Table S2. Vertices for *BHG*

| Feature | Resource Competition | Resource Defense | Mobility/Exchange |
|---|---|---|---|
| g2g1 | 6.071733 | 2.976645 | 2.45832 |
| prevalue | 16.24931 | 4.628441 | 5.590594 |
| huntfil2 | 1 | 1 | 1 |
| g2mhs | 16.7231 | 12.70297 | 4.749564 |
| mhs | 19.98424 | 3.707549 | 3.957995 |
| gath | 1 | 1 | 3 |
| nomov | -4.22284 | 15.65726 | 14.32136 |
| bodyt | 1 | 2 | 1 |
| slave | 3 | 1 | 1 |
| lkmov | -0.10433 | 2.322361 | 2.949305 |
| occspe | 4 | 1 | 1 |
| divmor | 1 | 3 | 1 |
| szmean | 14.17401 | 1.654321 | 2.436451 |
| g1mhs | 2.706253 | 4.354268 | 2.109871 |
| agedif | 3.536832 | 14.29632 | 2.963695 |
| revres | 1 | 3 | 1 |
| money | 4 | 1 | 1 |
| dkinex | 3 | 1 | 1 |
| lkspmov | 0.555912 | 1.132914 | 1.706228 |
| polygrecod | 5.418128 | 39.08245 | 0.034362 |
| levira | 2 | 2 | 1 |
| wx.polygny | 8.808877 | 45.37002 | -2.54708 |
| agecom | 1 | 4 | 1 |
| discomp2 | 1 | 2 | 1 |
| marrycer | 4 | 1 | 1 |
| initm | 2 | 5 | 0 |
| war1 | 4 | 1 | 1 |
| g2basord | 11 | 4 | 3 |
| perogat | 5 | 1 | 1 |
| agem | 18.72782 | 26.57441 | 17.78429 |
| sorora | 2 | 2 | 2 |
| enemy | 4 | 1 | 1 |
| grppat | 3 | 1 | 1 |
| initexm | 2 | 6 | 0 |
| polygn | 3 | 6 | 2 |
| boyseg38 | 1 | 3 | 1 |
| shaman | 4 | 2 | 1 |
| prison | 4 | 1 | 1 |
| store | 4 | 0 | 3 |
| disloc | 4 | 1 | 1 |
| divorce | 1 | 2 | 3 |
| fishing | 88.59382 | 4.865173 | 16.88931 |
| dspmov | 2.789693 | 8.593764 | 27.54367 |
| warlead | 3 | 2 | 2 |
| deadav | 2 | 2 | 1 |
| marprop | 4 | 1 | 1 |
| mdivlab | 68.63252 | 32.67882 | 72.93477 |
| death | 4 | 1 | 1 |
| conpos | 3 | 2 | 1 |
| dritual | 3 | 3 | 1 |
| ritscal | 4 | 3 | 1 |
| mardir | 2 | 2 | 2 |
| agef | 15.1915 | 12.27689 | 14.82008 |
| lpackinx | 1.503836 | 0.159899 | -1.00691 |
| gpgpcon | 4 | 2 | 1 |
| sed | 5 | 1 | 2 |
| fish | 3 | 3 | 4 |
| polyscal | 4 | 1 | 2 |
| qtstor | 4 | 0 | 3 |
| subdiv2 | 69.44157 | 76.64706 | 71.34456 |
| packing | 2 | 2 | 1 |
| kinexo | 4 | 3 | 1 |
| packord | 4 | 3 | 2 |
| minlaw | 1 | 3 | 1 |
| housex | 7 | 0 | 3 |
| intcon | 3 | 2 | 2 |

### Table S3. Features used from *Pulotu*

Table is attached as supplementary, and can be interpreted using the codebook available at https://pulotu.econ.mpg.de/.

### Table S4. Features used from *BHG*

Table is attached as supplementary, and can be interpreted using the codebook available at http://ajohnson.sites.truman.edu/data-and-program/.

### Table S5. Limit of data.

Table is attached as supplementary.

### Table S6. Traits associated with vertices for *Pulotu*

Table is attached as supplementary. Statistically significant associations are marked with either (*) for $p<0.02$ or (**) for $p<0.0002$. We describe the regression in Supplementary Information 6.

### Table S7. Traits associated with vertices for *BHG*

Table is attached as supplementary. Statistically significant associations are marked with either (*) for $p<0.02$ or (**) for $p<0.0002$. We describe the regression in Supplementary Information 6.

### Table S8. Distances from archetypes for *Pulotu*

Table is attached as supplementary. Note that distances are normalized using z-score.

### Table S9. Distances from archetypes for *BHG*

Table is attached as supplementary. Note that distances are normalized using z-score.

Table S10. Data for the relation between island type and distance from archetypes for Polynesian and Micronesian cultures.

| Culture | Island Type | Island Size (km^2) | Maximum Elevation (m) | Population | Mobility/Exchange | Resource Competition | Resource Defense |
|---|---|---|---|---|---|---|---|
| **Anuta** | Volcanic High Island | 0.4 | 78 | 125 | 7.9 | 3.6 | 8.9 |
| **Chamorro** | Volcanic High Island | 541 | 406 | 40000 | 4.1 | 8.3 | 6.6 |
| **Chuuk** | Volcanic High Island | 34.2 | 443 | 12000 | 7.7 | 3.6 | 9.1 |
| **Fiji (East)** | Tectonic | 10429 | 1324 | 90000 | 8.7 | 4.1 | 7.3 |
| **Futuna (East)** | Volcanic High Island | 46 | 760 | 1000 | 5.7 | 6 | 7.4 |
| **Futuna (West)** | Volcanic High Island | 10.7 | 589 | 1000 | 5.6 | 8 | 5.1 |
| **Hawaii** | Volcanic High Island | 10458 | 4206 | 250000 | 9.3 | 2 | 10 |
| **Kapingamarangi** | Atoll | 1.1 | 5 | 300 | 3.5 | 7.8 | 8.5 |
| **Kiribati** | Atoll | 49 | 4 | 35000 | 6.6 | 5.5 | 6.9 |
| **Mangaia** | Atoll | 52 | 169 | 2500 | 9.5 | 2.4 | 9 |
| **Mangareva** | Volcanic High Island | 15.4 | 441 | 1500 | 7.3 | 4.1 | 8.5 |
| **Manihiki and Rakahanga** | Atoll | 5.4 | 3 | 1200 | 5.8 | 5.5 | 8.5 |
| **Maori** | Continental island | 116219 | 2797 | 100000 | 9.6 | 2.3 | 9 |
| **Marquesas** | Volcanic High Island | 339 | 1185 | 35000 | 9.2 | 2.2 | 10 |
| **Marshall Islands** | Atoll | 16 | 5 | 9000 | 4.8 | 6.9 | 7.4 |
| **Moriori** | Volcanic High Island | 900 | 287 | 2000 | 4.9 | 6.9 | 7.1 |
| **Niue** | Atoll | 259 | 70 | 5000 | 7.6 | 4 | 8.2 |
| **Ontong Java** | Atoll | 8 | 2 | 2000 | 4.3 | 7 | 8.9 |
| **Palau** | Volcanic High Island | 370 | 217 | 25000 | 6.6 | 6.4 | 5.8 |
| **Pohnpei** | Volcanic High Island | 336.7 | 778 | 10000 | 6.6 | 4.8 | 8.7 |
| **Pukapuka** | Atoll | 1.3 | 30 | 500 | 4.8 | 6.5 | 8.6 |
| **Rapa Nui** | Volcanic High Island | 163 | 600 | 3000 | 9.8 | 2.8 | 8.4 |
| **Rarotonga** | Volcanic High Island | 67.1 | 653 | 7000 | 9.7 | 1.7 | 10 |
| **Rennell and Bellona** | Atoll | 840 | 180 | 1450 | 4.6 | 7 | 7.6 |
| **Samoa** | Volcanic High Island | 1091 | 1100 | 60000 | 9.7 | 3.4 | 7.9 |
| **Tahiti** | Volcanic High Island | 1040 | 2237 | 45000 | 9.1 | 2.6 | 9 |
| **Tikopia** | Volcanic High Island | 5.7 | 360 | 1000 | 7.9 | 3.5 | 9.1 |
| **Tokelau** | Atoll | 4 | 3 | 750 | 5.4 | 6.2 | 7.6 |
| **Tonga** | Tectonic | 260 | 70 | 35000 | 7.5 | 4 | 8.4 |
| **Ulithi** | Atoll | 4.5 | 6 | 800 | 4.2 | 7.6 | 7.2 |
| **Woleai (Ifaluk)** | Atoll | 1.5 | 6 | 300 | 3.7 | 7.6 | 8.8 |
| **Yap** | Volcanic High Island | 100.2 | 178 | 8000 | 3.9 | 8.6 | 6.5 |
